# The Preprint Club - A cross-institutional, community-based approach to peer reviewing

**DOI:** 10.1101/2023.01.04.522570

**Authors:** Felix Clemens Richter, Ester Gea-Mallorquí, Nicolas Ruffin, Nicolas Vabret

## Abstract

The academic community has been increasingly using preprints to disseminate their latest research findings quickly and openly. This early and open access of non-peer reviewed research warrants new means from the scientific community to efficiently assess and provide feedback to preprints. Yet, most peer review of scientific studies performed today are still managed by journals, each having their own peer review policy and transparency. However, approaches to uncouple the peer review process from journal publication are emerging. Additionally, formal education of early career researchers (ECRs) in peer reviewing is rarely available, hampering the quality of peer review feedback. Here, we introduce the Preprint Club, a cross-institutional, community-based approach to peer reviewing, founded by ECRs from the University of Oxford, Karolinska Institutet and Icahn School of Medicine at Mount Sinai. Over the past two years and using the collaborative setting of the Preprint Club, we have been discussing, assessing, and providing feedback on recent preprints in the field of immunology. In this article, we provide a blueprint of the Preprint Club basic structure, demonstrate its effectiveness, and detail the lessons we learned on its impact on peer review training and preprint author’s perception.

## Introduction

Preprints are openly accessible and, thus, democratise the dissemination of scientific information. Their public availability invites a large range of feedback from the research community allowing for a broader assessment of research findings. Fields such as mathematics and physics have been embracing the use of preprints for several decades. However, the biomedical research field, including the field of immunology, remained hesitant on disseminating their research prior to publication in peer reviewed journals, despite concerted efforts to promote their use^1,2^. Nonetheless, the deposition of non-peer reviewed life science manuscripts to preprint servers has grown nearly exponentially over the past years^3^. Its adoption by immunologists and other communities from related field has dramatically increased during the COVID-19 pandemic, which resulted in an expedited exchange of knowledge and accelerated the scientific response to this outbreak^4^. Preprint servers became a primary access point of information used by researchers to develop and contextualise emerging COVID-19 data, with an estimated 25% of all COVID-19–related publications deposited on preprints servers between January and October 2020^5^. This highlights that the benefits of preprints for fast, accessible research distribution have become clear to the wider biomedical research community. The flood of COVID-19 related preprint depositions, which was estimated at 39.5 preprints per day during the first months of the pandemic^4^, made it difficult for many frontline researchers to filter relevant and rigorous studies and to stay ahead of the information wave. In addition, once released, preprints quickly taken up by media outlets may have caused cases of misinformation^6^. Consequently, the review of preprints is an emerging need arising from changes in publication practice.

Among the various approaches proposed in preprint peer review^7^, a critical role can be played by community-based assessments, such as journal clubs, as they allow providing feedback on scientific findings after discussion among peers with diverse background and may prove useful for training critical thinking among early career researchers (ECRs)^8^. Thus, during the COVID-19 pandemic, ECRs from the immunology departments of the Icahn School of Medicine at Mount Sinai and the University of Oxford have taken advantage of their journal club communities to write and share public reviews of selected preprints and highlight important studies that would help on the understanding of the immune response to SARS-CoV-2^9,10^.

Using this experience, in November 2020, we founded the Preprint Club, an initiative using a community-based approach to evaluating and digesting immunology-focussed preprints. Together, PhD students, postdoctoral fellows and faculty members form a ‘Preprint Club hub’ to discuss, assess and give feedback to preprints on a weekly basis in this cross-institutional journal club setting. In each session, two ECRs (either PhD students or postdoctoral fellows) from alternating institutions present preprints that they consider interesting to the community at large. After each presentation, the members of the Preprint Club hub anonymously score the presented preprints based on their *Novelty, Significance* and *Scientific Quality*. All the presenters are encouraged to write up a short review on the preprint they presented, which are posted as a public review repository (www.preprintclub.com) and linked to the original preprint to inform authors about the feedback. Through a collaboration with *Nature Reviews Immunology* (NRI), the best preprints that passed community-based threshold scores and criticism, are selected to be further highlighted this journal. In addition, we use social media (i.e. Twitter: @preprintclub) to communicate the peer reviews to the preprint authors, to other immunologists and the wider scientific community. A detailed depiction of the Preprint Club, its system and structure, is provided in **Fig. 1**.

**Figure 1:**
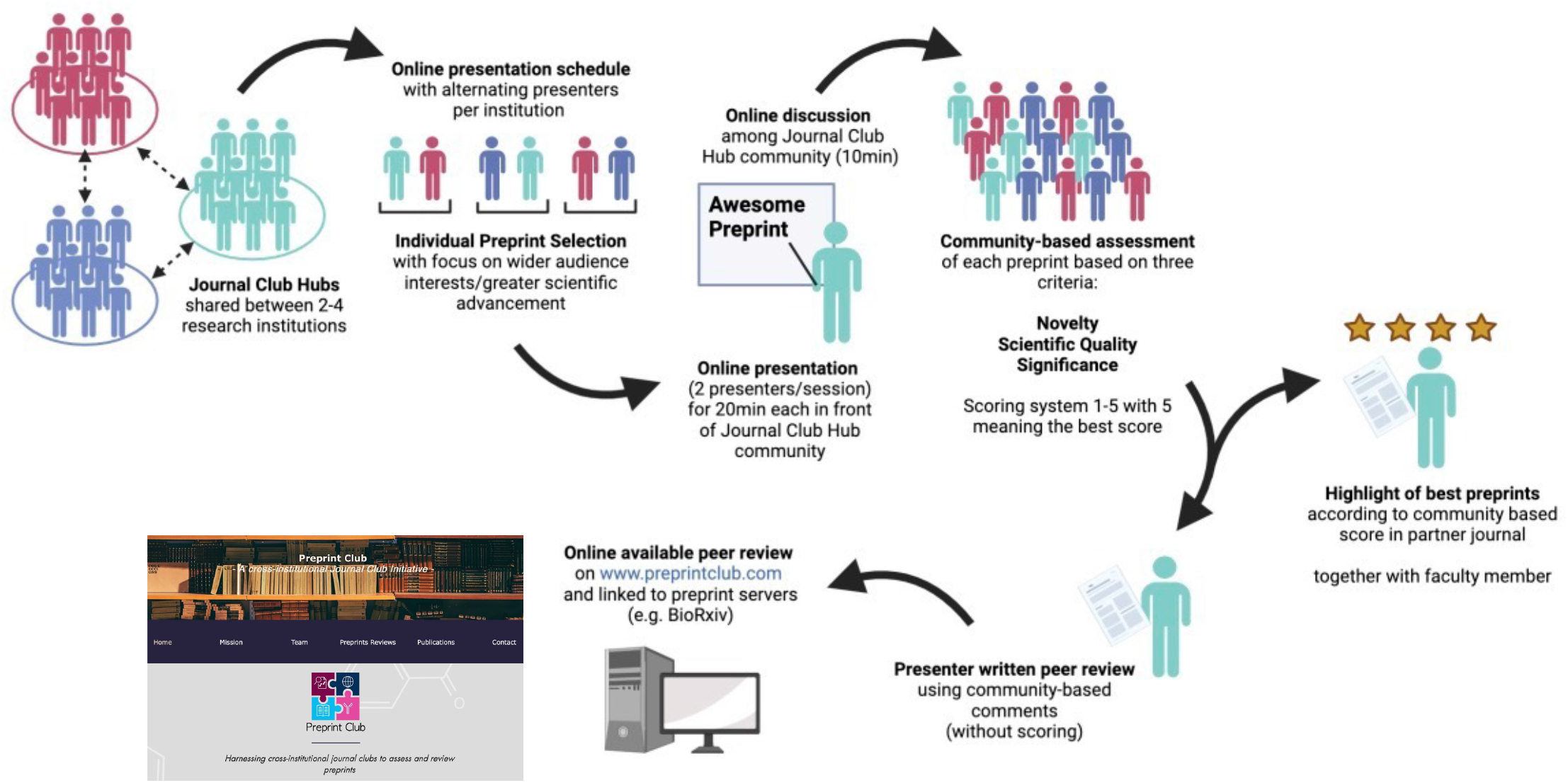
Organizational structure of the Preprint Club. The Preprint Club is organized in small hubs of 2-4 collaborative institutions. Once a Preprint Club hub has formed, an online schedule is formed with alternate presenters, ideally making sure that presentation slots are evenly distributed. Two presenters are scheduled per week and session. The presenters select a recent preprint of their choice, which is then presented and discussed during the online journal club session (20min + 10 min). Each participant uses a three-category voting system to rank the preprint with a score of 1 being the lowest and 5 being the highest. Preprints are assessed based on their novelty (are the findings novel and original?), their scientific quality (quality of experimental design and data obtained, and how well they support the authors’ conclusions) and their significance (how likely this study is going to impact its research field and immunology in general). After the journal club, presenters are encouraged to take aboard the feedback from the discussion and write up a peer review digest to give feedback to the preprint authors. This feedback is uploaded on the Preprint Club Website (www.preprintclub.com) and made publicly available. Each month, scores for all presented preprints are collected (~8 preprints/month). Preprints reaching a threshold (in our experience: ~11.5 or higher) are considered for further highlight in a partner journal (in our case: Nature Reviews Immunology). The highlight on the preprint is written together with a faculty member from the partnering institutions to improve the conclusions drawn from the preprint assessment. This system can be applied to many other biomedical research fields in the same way and require only motivated ECRs from different institutions, a partnering journal and an online meeting platform (e.g. Zoom, Teams).

While the Preprint Club system provides a novel way of community-based approach to peer review and an educational element to ECR’s training, little is known about the efficacy of such systems and how it is perceived by its participants and preprint authors. For this reason, we describe here the lessons learned from our experience alongside an evaluation of its predictive value using commonly used impact metrics such as journal impact factors (JIFs) and citation rates. We also performed two independent, anonymous surveys to learn more about its perception, training purposes and usability for Preprint Club presenters and preprint authors. Our initial data support the notion that (1): our community-based approach is able to select the most relevant work from non-peer reviewed data; (2): this assessment has training and educational value for a ECRs peer reviewing and (3): the reviews are generally well perceived and found helpful by preprint authors, altogether highlighting the potential benefit of community-based reviews to improve science publication practices.

## Results

### The Preprint Club assessment system allows selection of preprints with higher citation rate

From November 2020 to July 2022 (date of our data collection), the Preprint Club hub has discussed and assessed 113 preprints across 62 journal club sessions. Out of the 113 preprints, 60 (53%) have been published in a peer reviewed journal as of August 2022. From these, 24 preprints have been selected and highlighted in NRI. The selection system is based on the overall score given by the Journal Club participants that assesses the *Scientific Quality*, *Novelty*, and *Significance* of the presented preprint. After an initial period when the two best scored preprints every month were highlighted in NRI, we defined a threshold of 11.5 out of 15 of the overall preprint score to limit the highlights to top-quality preprints. Overall scores followed a gaussian distribution (R^2^= 0.89) and setting a threshold at 11.5 led to 11.5% of preprints to be highlighted in NRI (**Fig. 2A**).

**Figure 2:**
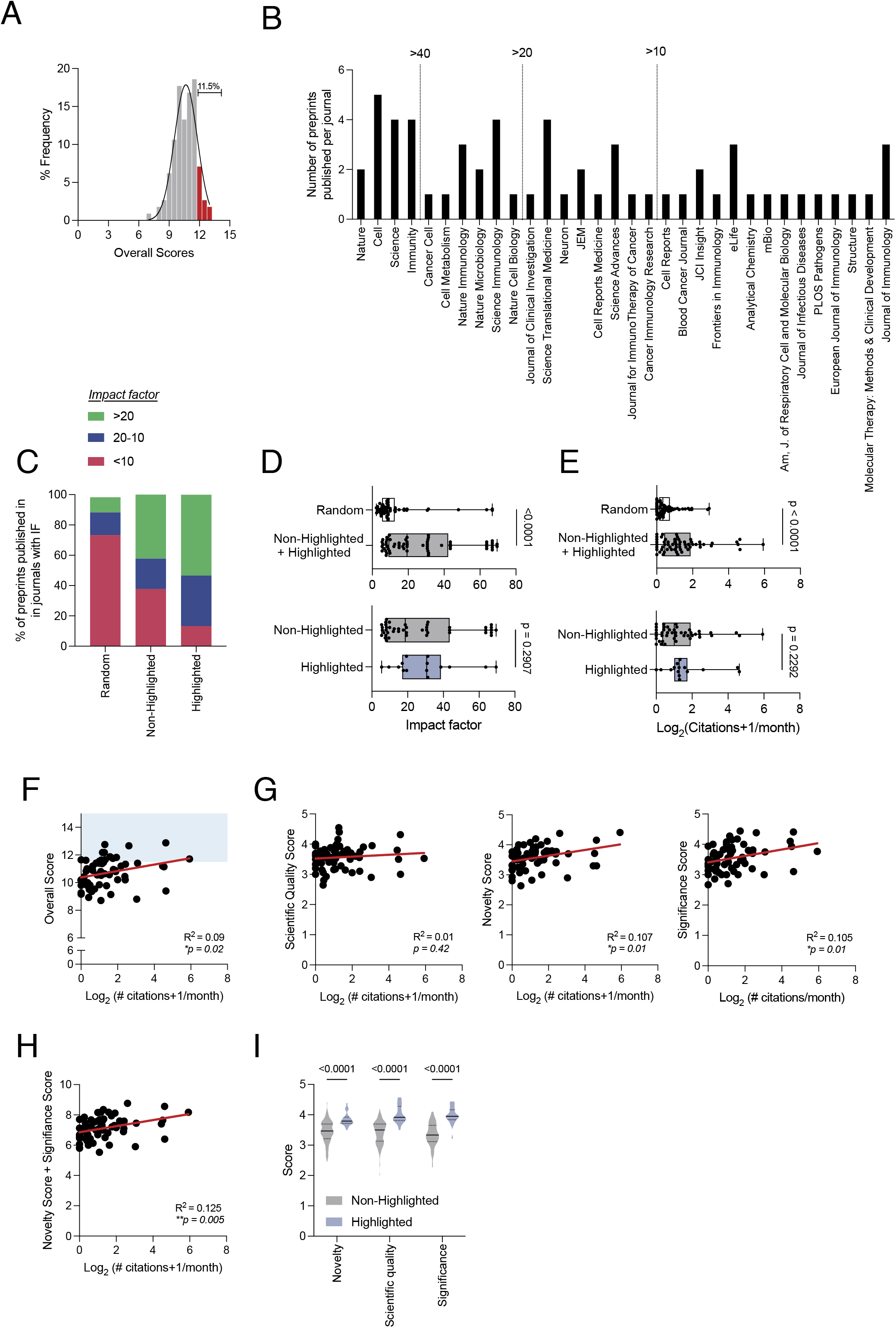
The Preprint Club assessment system enriches for preprints eventually published in higher impact factor journals and with higher citation rates. **(A)** Scoring of preprints follows a normal distribution pattern with the top 11.5%of preprints having a score of >11.5, prompting them to be considered for highlighting. **(B)** Journals where the preprints we assessed were eventually published in. **(C)** Distribution of preprints published in journals according to their corresponding impact factors. **(D+E)** Journal impact factor (**D**) and citation rate (**E**) of non-highlighted and highlighted preprints discussed in the Preprint Club compared with of a randomly selected group of immunology-related preprints (random). Mann-Whitney test. **(F)** Correlation of overall assessment scores with article citation rate. Blue shaded area marks preprints with an overall score higher than 11.5. **(G)** Correlation of each individual assessment score with paper citation rate. **(H)** Correlation of the combined score of novelty and significance with citation rate. **(I)** Comparison of each score between highlighted and non-highlighted preprints. Two-Way ANOVA with Sidak correction.

To measure the performance of the Preprint Club setup, we sought to determine the outcomes of preprints assessed during the Preprint Club sessions. We found that the preprints discussed have been eventually published in a broad spectrum of journals and publishers (**Fig. 2B**), indicating that their assessment did not *per se* poise the preprint to be published in specific journals. Next, we analysed whether and how the Preprint Club selection was enriched for preprints with a potential of a greater impact. We included in the analysis a control dataset made of randomly selected immunology preprints deposited on BioRxiv (group called ‘random’) during the whole month of September 2021. Compared to this ‘random’ group, we found that the pool of preprints chosen by the Preprint Club community were published in journals with higher JIF (**Fig. 2C**). We term this an “individual-based” enrichment effect, as the initial selection of the preprints that are presented in the Preprint Club depends strictly on the choice of the presenter. Altogether this shows the ability of individual ECRs to select relevant work. Furthermore, the dataset suggests that the preprints highlighted in *NRI* tended to be published in even higher impact factor journals (**Fig. 2C**). We term this a “community-based” enrichment effect, as the selection for highlights is based on the overall score attributed by the Preprint Club hub and highlights the ability of community-based assessment to select significant findings in our field.

JIFs are a controversial measurement of scientific impact that do not necessarily reflect the significance of published work^11,12^. Thus, we analysed the dataset with respect to both the JIF and the citation rate for each publication. Non-highlighted and highlighted preprints had significantly better citation rates and JIFs than the random preprint pool (**Fig. 2D-E**). Moreover, we found that the mean JIF and citation rate were higher in highlighted compared to non-highlighted preprint (**Fig. 2D-E**). Next, we correlated our community-based scoring system with the established parameters, both the citation rate and JIF of the published article. The overall preprint score modestly, but significantly, correlated with the citation rate (**Fig. 2F**). When stratified by each individual scoring category (*Novelty*, *Significance* and *Scientific Quality*), we found that both *Novelty* as well as *Significance* fitted best the citation rate, while *Scientific Quality* showed no correlation (**Fig. 2G**). Using this information, we found that combining the scores of *Novelty* and *Significance* can improve the preprint predictive value with regards to the citation rate (**Fig. 2H**). A similar analysis was also performed for the JIF but showed less or no correlation. To understand why the scoring category *Scientific Quality* did not seem to predict the outcome of the preprint, we compared the scores of highlighted and non-highlighted preprints and found that all categories were significantly higher in highlighted preprints (**Fig. 2I**). This suggests that while *Scientific Quality* is a scoring metric that allowed us to discriminate between preprints, it plays a minor role compared to *Significance* and *Novelty* in predicting future citation rate. Overall, we found that the Preprint Club assessments are in line with and can predict the interest of scientific community toward the specific preprints discussed, enriching for preprints with potentially great impact.

### Inclusion of a third institution ameliorates scoring biases

We next sought to identify potential biases involved both in the Preprint Club assessment system and the preprint selection choice. We first evaluated whether the affiliation of the preprint presenter has an impact on the overall score. Plainly, we analysed whether individuals from the same institution as the presenter would assess the preprint differently and bias the vote. We classified scores given by individuals from the same institution as the presenter (‘presenter’) and scores given by individuals from another institution (‘non-presenter’). Strikingly, we found that there was a voting bias during the initial period of the Preprint Club, when participants came from only two institutions. Scores were higher when given to preprint presenters from the same institution as compared to scores given to preprint presenters from another institution (**Fig. 3A, left**). After inclusion of a third institution, this bias no longer persisted (**Fig. 3A, right**). We further compared the scores given by members from each institution to understand whether specific institutions would score differently. Interestingly, we found that preprints received slightly lower scores from members of institution 2 (**Fig. 3B**). On the other hand, institution 1 and 3 scored similarly, independent of the presenter’s affiliation. Next, we wanted to understand whether specific institutions are better at picking preprints compared to other institutions. For this, the dataset was stratified by the presenter’s institution and by the overall assessment scores given to the preprint by all the Preprint Cub attendants. Institution 2 appeared to pick preprints with a slightly higher overall score, but the overall scores did not dramatically differ between institutions (**Fig. 3C**). Of note, less preprints were presented by institution 3, which could explain the observed differences. We hypothesised that these differences would also translate into better JIF and citation rate outcomes. Indeed, we found that preprints presented by institution 2 had the highest citation rate and JIF (**Fig. 3D**). Taken together, we found limited biases in the Preprint Club assessment system which can largely be overcome through inclusion of three or more institutions to balance differences in scoring behaviours.

**Figure 3:**
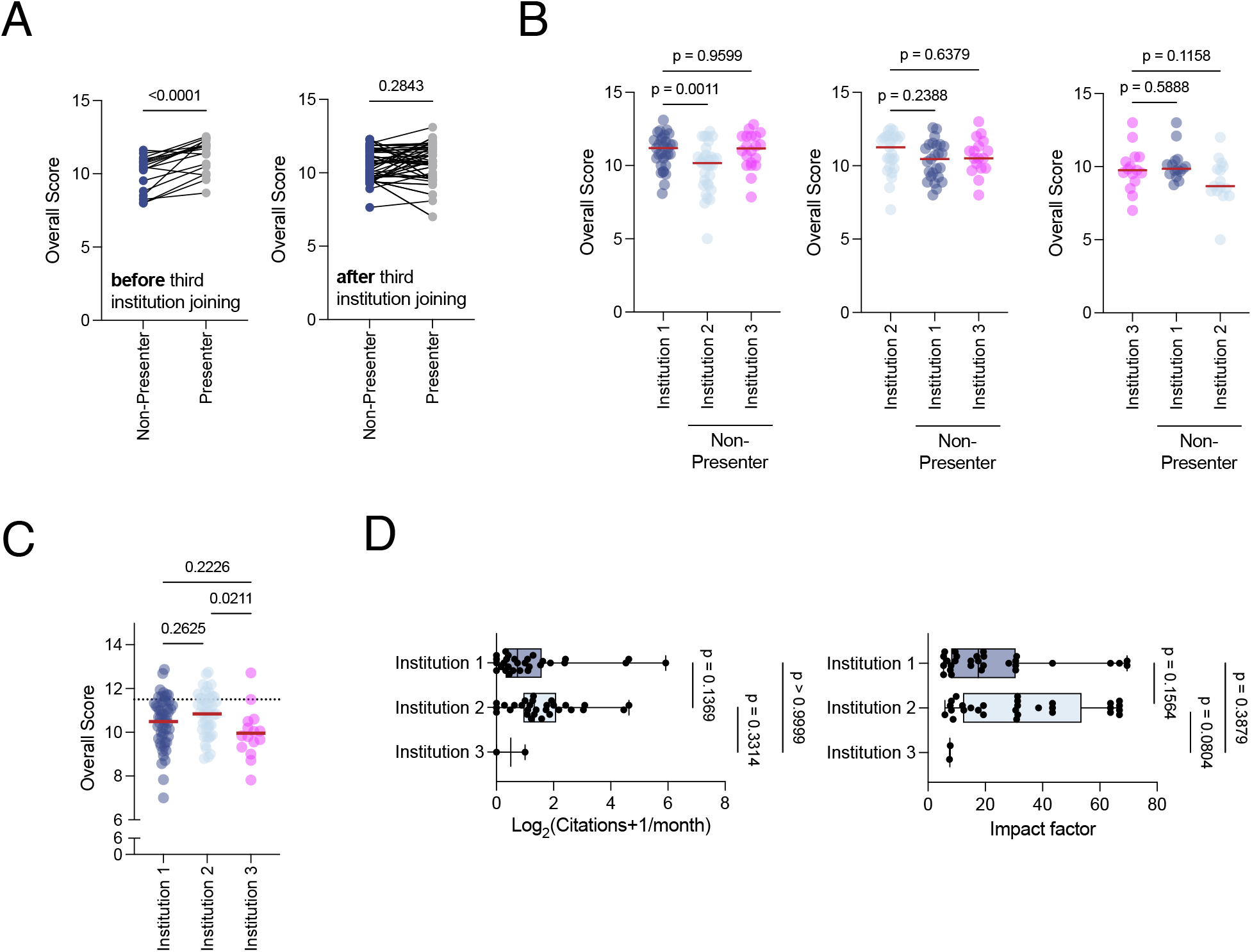
Bias of the Preprint Club assessment system. **(A)** Assessment of overall scores given from individuals of the same institution as the presenter (‘Presenter’) in comparison to individuals who do not belong to the same institution than the presenter (‘Non-Presenter’), before and after addition of a third institution in the Preprint Club. Paired t-test. **(B)** Overall score per institution by presenting institution. One-Way ANOVA. **(C)** Overall score received from different institutions. One-Way ANOVA. **(D)** Outcomes of preprints selected by institution. Kruskal-Wallis test.

### Preprint Club assessments are well perceived by preprint authors

Although the Preprint Club assessment system performed well in identifying preprints with high potential impact, we were less certain about the perception of unsolicited reviews by the preprint authors themselves. To address this question, we sent an anonymous survey to the first and corresponding authors of the preprints presented to ask for their opinion about the Preprint Club’s assessment of their work. The survey’s objectives were to (1) learn if and how the authors were familiar with the preprint club, (2) collect feedback about whether the Preprint Club’s review of their study was found useful and (3), when applicable, collect feedback about the corresponding highlight published in *Nature Reviews Immunology*. Details of the survey and authors’ replies are included in **supplementary table 1**.

The results show that while most of the authors knew about the Preprint Club, the majority learned about it after we reviewed their preprint (**Fig. 4A**). ECRs and junior principal investigators (PIs) were more likely to know about the preprint club compared to senior PIs (**Fig. 4A**). While most authors had learnt about the Preprint Club via Twitter, about 30% of all respondents got to know the initiative through the publication of a highlight in NRI or through a widget embedded on preprint BioRxiv pages that notifies preprint authors that a preprint has been the subject of a *Community Review* (**Fig. 4B**). In addition, authors of highlighted preprints were more likely to know that their preprint had been reviewed (87%) compared to authors of non-highlighted preprints (58%). Although unsurprising, this emphasises the benefits of partnerships of community reviews with scientific journals and editors to increase the visibility of similar initiatives.

**Figure 4:**
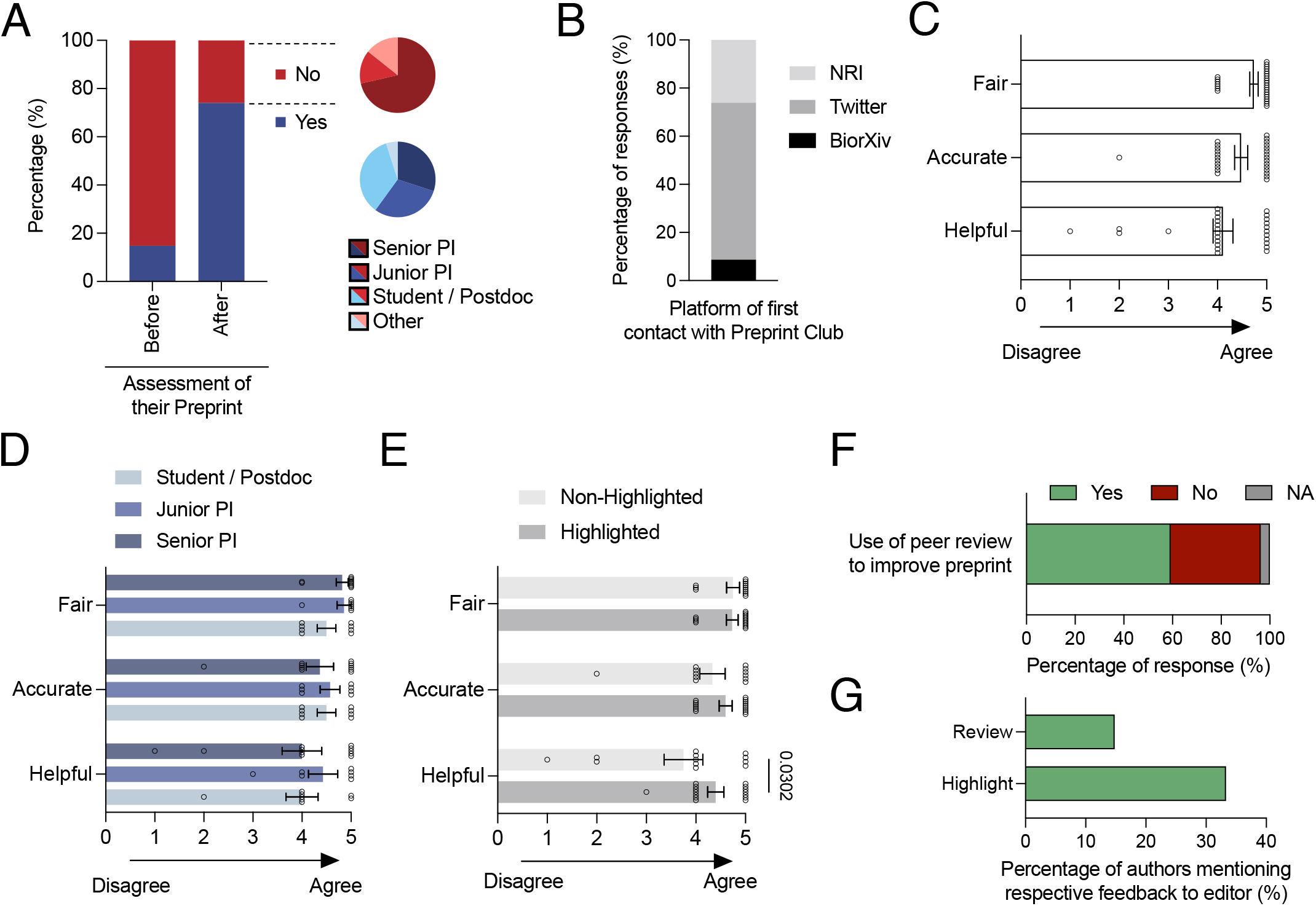
Authors’ opinion of their preprint assessment by the Preprint Club. **(A)** Fraction of preprint authors aware of the existence of the Preprint Club either before or after their preprint was reviewed. **(B)** Sources first used by preprint authors to learn about Preprint Club. **(C-E)** Scores given by preprint authors to the review written on their preprint (**C**), broken down by authors status (**D**) or whether the preprint was further highlighted or not (**E**). Two-Way ANOVA. **(F)** Fraction of authors who used the content of reviews to improve their manuscript. **(G)** Fraction of authors that mentioned the review or the highlight in subsequent communication with journals editors. ***(A-C)** Replies from PhD students and Postdoc were grouped for readability.*

To get a better understanding of the quality of the review written by Preprint Club members, we asked preprint authors to evaluate the *helpfulness, accuracy*, and *fairness* of our review. Overall, preprint authors rated the reviews positively, with respondents scoring *fairness* highest, followed by *accuracy* and *helpfulness* (**Fig. 4C**). The perception of the Preprint Club reviews was independent of the career status of the respondents (**Fig. 4D**). Importantly, when the responses were stratified on whether the preprint was highlighted or not, little differences were observed, with only “helpfulness” tending to be higher (0.65 points) in highlighted preprints (**Fig. 4E**). This may be explained by the fact that preprint reviewers may write a more detailed preprint assessment when knowing the preprint will also be highlighted. Importantly, 59% of preprint authors used the feedback from the Preprint Club reviews to improve their manuscript (**Fig. 4F**) but only a minority mentioned the review or the highlights (14.8% and 33.3%, respectively) when further communicating with journal editors (**Fig. 4G**).

Finally, when asked for feedback on how to improve the Preprint Club, or which feature authors would like to see added, the replies were overwhelmingly positive (**Suppl. Table 1**). Several authors indicated that they would prefer a direct notification once a review has been posted or to join the Preprint Club session that discussed their preprint. Another author suggested having the option for authors to reply to the review on the Preprint Club website.

Overall, this survey showed preprint authors had a generally positive perception of the assessment of their work by the Preprint Club, independent of the fact that their preprints were further highlighted in NRI or not. This encouraging feedback, collected anonymously, is of importance as it could alleviate concerns about the preprint author’s perception of similar initiatives.

### Community-based reviewing systems are well perceived and preferred to ‘classical’ journal-dependent peer reviewing processes

Aside from measuring the performance of Preprint Club, we wanted to understand the mechanisms behind the selection and identification of preprints by the Preprint Club community. We found that most presenters in the Preprint Club identified a preprint of interest by screening through preprint servers, with BioRxiv being the most popular (**Fig. 5A**). However, about 20% of all presented preprints were either found via Twitter or were recommended by a colleague. Overall, most people screened between 5-10 preprints before selecting a preprint for presentation at the Preprint Club (**Fig. 5B**). Interestingly, most Preprint Club participants focused their search hereby on the abstract and the figures of the preprint, while only a minority screened the full paper (**Fig. 5C**). Preprint Club participants focused on a good balance of *Novelty*, *Scientific Quality*, and *Significance*, with the latter being the single most important criteria of selection (**Fig. 5D**). The use of the *Significance* of the finding for the research field to select the preprints may also explain its overall increased correlative value in identifying preprints with higher citation rates (**Fig. 2G**). Importantly, most presenters selected a preprint from a group unknown to them (58.6%) indicating that a limited bias in selection process by the status of given research groups. Taken together, this data highlights that participants in the Preprint Club select preprints typically by screening a limited number of preprints with the highest overall balance of selection criteria.

**Figure 5:**
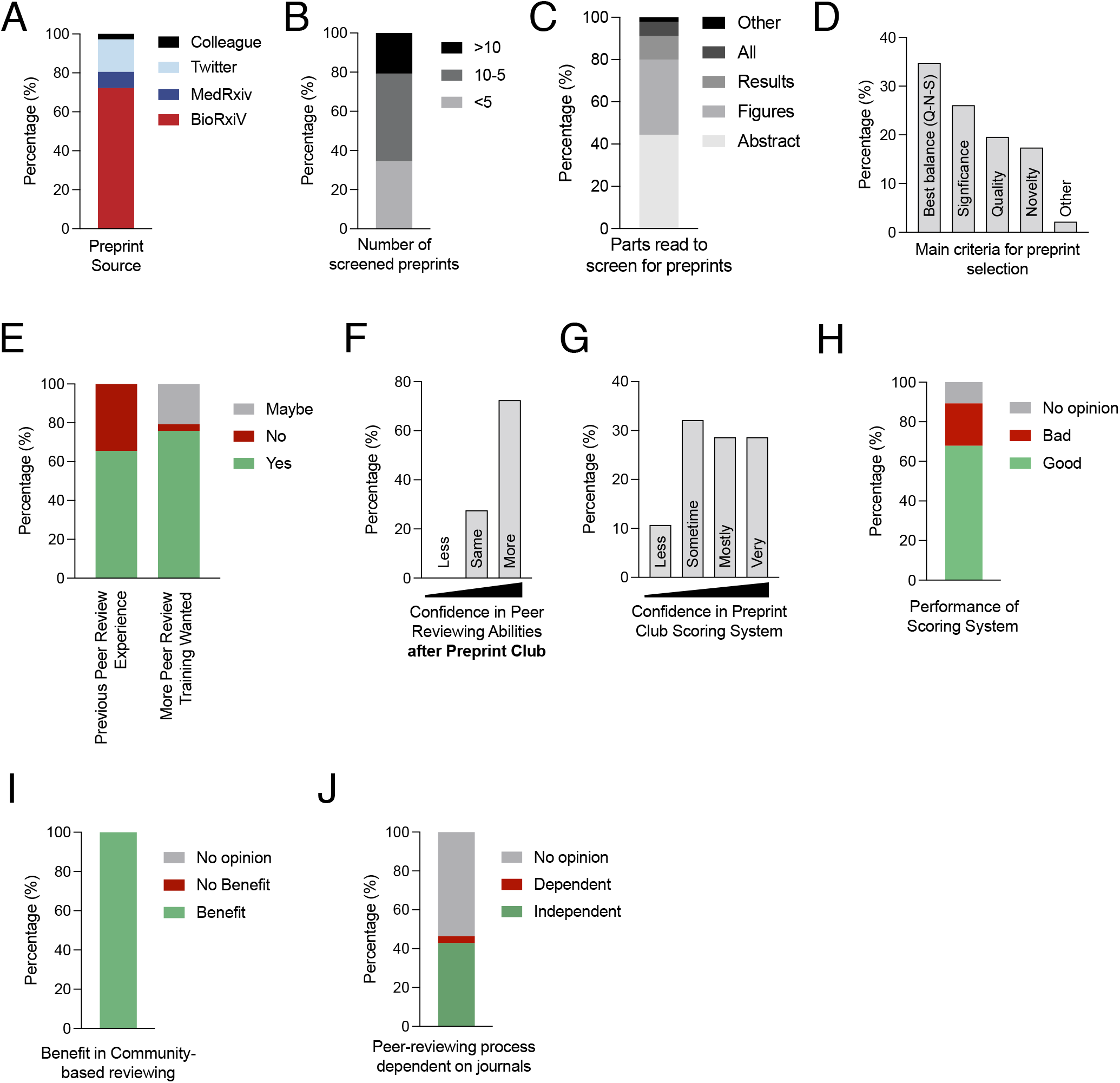
Community-based reviewing systems are well perceived by members of the Preprint Club and preferred to ‘classical’ journal-dependent peer reviewing processes. **(A)** Sources used by presenters to identify preprints of interest. **(B)** Numbers of preprints assessed before choosing a preprint for presentation. **(C)** Preprints sections initially assessed before choosing a preprint for presentation. **(D)** Main criteria used to select preprints. **(E)** Prior experience in peer reviewing before taking part in the Preprint Club and request for training. **(F)** Confidence in peer review after taking part in the Preprint Club. **(G)** Confidence in the Preprint Community’s ability to assess a given preprint. **(H)** Perception of the performance of the scoring system. **(I)** Fraction of Preprint Club members seeing a benefit in community-based reviewing. **(J)** Fraction of Preprint Club members that believe the peer reviewing process should depend on the journal.

### Community-based review improves ECR’s confidence in peer review skills

Next, we evaluated the training value of the Preprint Club. Remarkably, over 50% of all participants had previous peer review experience, but more than 75% considered they would benefit from a formal peer review training (**Fig. 5E**). Since one of the goals set by the Preprint Club was to train ECRs in delivering understandable and constructive peer reviews, we sought to determine whether participants in the Preprint Club would feel more confident in evaluating and reviewing a manuscript if asked for. More than 70% felt that the participation in the Preprint Club made them more confident in assessing scientific research manuscripts (**Fig. 5F**). This data underlines the need for more formal peer review training and shows the benefits of community-based discussions like the preprint-based journal club initiatives, which focus on yet unpublished research and may give more space for discussion. We also asked about the participant’s perception of the Preprint Club scoring system. The majority of participants felt very or mostly confident in assessing the preprints presented using the current scoring system (**Fig. 5G)** and had an overall positive perception of its design (**Fig. 5H**).

More recently initiatives such as Reviews Commons, which allows for a journal-independent peer reviewing process, have become increasingly popular. We were therefore interested how the Preprint Club community perceived different peer review processes. Perhaps unsurprisingly, participants in the Preprint Club found a clear benefit in a community-based reviewing system (**Fig. 5I**). Furthermore, 42% thought that peer review should be uncoupled from the journals (**Fig. 5J**). About 50% of the Preprint Club participants did not have a particular opinion whether one system is better than the other and only a small fraction (3.5%) would certainly want to maintain the current peer review system. This feedback suggests that there is a potential for more community-based, journal-independent peer reviewing processes in academic publishing.

## Discussion

Preprint peer review is a growing trend in the scientific community, where researchers can share their work in a preprint server and collect feedback from their peers before submitting their manuscript to a journal for formal peer review. This approach has several benefits, including faster dissemination of research findings, greater transparency in the review process, and the opportunity for researchers to receive early feedback on their work.

Several preprint peer review initiatives have emerged over the recent years, providing different services, such preprint reviewing platforms (e.g. Rapid Reviews, Peer Community In, PREreview) or highlighting preprints to the broader research community (e.g. Prelights). A working group led by the preprint-focussed research organisation Accelerating Science and Publication in Biology (ASAPbio) recently defined 8 key features of the preprint peer review process to allow for a better classification of different preprint peer review initiatives and their functions^7^. According to this classification, in the Preprint Club, (i) the reviews are requested by non-authors, (ii) the reviewer is self-nominated among our community of ECRs from partnering institutions, (iii) public’s interaction is included, (iv) there is no inclusion of author’s response, (v) no decision is provided at the end of the review process, although a community score (not publicly disclosed) allows to recommend preprints for subsequent highlight, (vi) the review covers the entire paper, (vii) the choice of being anonymous is at the reviewer’s discretion, although all our reviewers have decided to sign their reviews so far, and (viii) no competing interest statement is provided but an internal check makes sure we do not review preprints from our partnering institutions.

In 2019, the cross-journal initiative Reviews Commons, led by EMBO press, allowed for a journal-independent reviewing process where authors receive feedback on their preprint prior to their submission to associated journal’s editorial boards^13^. The boards then decide if the manuscripts enter a formal revision process and may eventually publish their article. From January 2023, the journal *eLife* is taking another direction. After initial editorial assessment, *eLife* now publishes every preprint sent for peer review as a “Reviewed Preprint” that includes an *eLife* assessment, public reviews, and a response from the authors, when available. The idea is to evolve from a classical ‘accept-or-reject’ bimodal decision after peer review, to provide instead a reviewer-based assessment of research findings, with the option of version-controlled improvement of the manuscript. Using this approach, the publisher hopes to “combine the immediacy and openness of preprints with the scrutiny of peer review by experts”^14,15^. This move of a well-renown publisher sparked a larger debate on the necessity of peer review, the value of journal names and the requirement of fast-pace publishing without compromising quality^16^.

The Preprint Club rallies around two major ideas: First, we wish to harness the force of institutional journal clubs to discuss, assess and peer review new research findings, and second, we aim at providing ECRs training and a first-hand experience in the peer review process. Indeed, journal clubs are educational entities that already exist in most institutions. They generally focus on most recent literature and harness knowledge that could provide feedback to authors of the articles they discuss^17^. Since journal clubs are also a place where students learned how to critically evaluate scientific manuscripts, they provide an ideal setting to learn the peer review process.

Despite advances proposed by *eLife* and other preprint review platforms, the peer review process remains largely organized and controlled by publishers. Shifting peer review away from journals offers several advantages, including lowering demand for redundant peer review through submission to multiple journals. Since peer review is a necessary part of academic publishing, well-trained peer reviewers remain essential for this process to ensure high research quality, improve scientific findings and advance the research field. Publishing groups (such as *Springer* or *Elsevier*) and independent organisations (such as *PREreview*) offer online peer review courses. However, for most ECRs, peer review training today primarily relies on interactions between trainees and their supervisors^18^. In this context, the Preprint Club provides a novel approach to train ECRs in peer reviewing and our data shows that with their experience in the Preprint Club, ECR became increasingly confident in their peer review skills.

Aside from holding an educational value to ECR, the initial data collected over the first 18 months of the Preprint Club highlights the ability of the Preprint Club to identify impactful manuscripts. However, a continuous assessment of the metrics used to measure its efficacy will be required to examine its longterm validity. Our initial performance analysis of the Preprint Club system revealed a better correlation of the preprint score with its citation rate than the impact factor of the journal they were published in. This may underline the shortcoming of JIFs as an indicator of the quality of publications, as debated previously^11,12,19^.

We found that the current setting can overcome a series of biases in assessing preprints, although a possible key confounding factor of the score given to the preprint might be the performance of the presenter itself. It appears plausible that the better a preprint is presented, the better it may be scored by the Preprint Club community as it may increase its understanding from the audience. Assessment of this bias may be helpful to perform more accurate scoring in the future.

Finally, while not solicited by preprint authors, our data indicate that the reviews from the Preprint Club were nonetheless well-perceived and their fairness, accuracy, and helpfulness were positively evaluated. Specific comments collected from the authors indicate that making sure the reviews are accessible before their manuscript is sent to traditional peer review could further improve the helpfulness as it would provide more time to incorporate edits in future versions of the manuscript.

Overall, our findings suggest that community-based peer review of preprints following the model of the Preprint Club is easily implementable in academic institutions and can provide a valuable addition to the traditional peer review process, offering benefits such as faster assessment of research findings and early training to peer review. For this reason, we encourage other academic institutions interested in the Preprint Club model to use the blueprint provided in this article to setup their own Preprint Club hubs with partnering institutions. We, the organisers of the Preprint Club, are happy to help you in setting up a similar system for your field, advise you and host your preprint digests on our collective website. The Preprint Club, so we are convinced, provides a novel element to publishing, scientific education and exchange across research institutions that can improve scientific communication and research culture.

## Methods

### Assessment of Preprint Club Effectiveness

All preprints that were presented in the context of the Preprint Club the October 2020 to June 2022, were used for the assessment of the effectiveness. We searched for the corresponding published articles, the journal they were published in, their JIF and their number of citations since publishing in the journal. The impact factors were from 2021 according to: https://impactfactorforiournal.com/icr-2021/, for impact factors that were not included in this list, an online google search was performed to find the JCR-2021 impact factor. Citations up to the 1^st^ August 2022 were included in the analysis. For the calculation of the citation rate, we removed one citation for each highlighted preprint to remove our own NRI highlight from overall citation rate. We included all preprints deposited in September 2021 on BioRxiv Immunology as a ‘random’ control group.

### Online Surveys

All online surveys for preprint authors and members of the Preprint Club community were performed anonymously using the Google Forms. The content of each survey and the replies we received are listed in Supplementary Table 1 and 2. The first survey was sent to the corresponding authors of each preprint that we reviewed during the period of interest, and we asked the corresponding author to forward the survey to the first author(s) of the preprint, when different (**Supplementary Table 1**). The second survey was sent to all the Preprint club members who presented one or more preprints during a journal club session (**Supplementary Table 2**).

### Statistics

Data were tested for normality before applying parametric or non-parametric testing. For correlation analysis, we performed a simple linear regression. Data was visualized and statistics calculated in either GraphPad Prism 9.

## Acknowledgments

The authors would like to thank to all the members of the Preprint Club for their hard work and dedication. This initiative would not have been possible without the support of the hosting institutions and the support of all involved faculty members. We would like to acknowledge all reviewers from the University of Oxford, the Icahn school of Medicine at Mount Sinai and Karolinska Institutet: Aljawharah Alrubayyi, Ghada Alsaleh, David Arcia-Anaya, Nima Assad, Austeja Baleviciute, Dorothée Berthold, Elena Brenna, Rachel Bond, Mariana Borsa, Graham Britton, India Brough, Matthew Brown, Yonina Bykov, Eduardo I. Cardenas, Lauren Chang, Bethany Charlton, Alice Chen-Liaw, Ewoud Compeer, Fabian Fischer, Micon Garvilles, Oksana Goroshchuk, John Hamp, Wanlin He, Samarth Hegde, Lisha Jeena, Rebecca Jeffery, Zerina Kurtovic, Vivian Lau, Matthew Lin, Stephanie Longet, Gabrielle Lubitz, Thi My Hanh Luong, Jaime Mateus-Tique, Julie M. Mazet, Guillaume Mestrallets, Chang Moon, Susanne Neumann, Jenna Newman, Tim O’Donnell, Matilde Oviedo Querejazu, Laura Palma Medina, Matthew Park, Isabela Pedroza-Pacheco, Luisanna Pia, Rachel Plitt, Tamar Plitt, Max Quastel, Rahul Ravindran, Ashley Reid, Mezida Saeed, Miriam Saffern, Christos Sazeides, Barbora Schonfeldova, Joan Shang, Prerna Suri, Michelle Tran, Abishek Vaidya, Verena van der Heide, Eveline van Gompel, Natalie Vaninov and Yavuz Yazicioglu. We would like to thank the preprint authors and reviewers who answer our anonymous surveys. Additionally, we appreciate all the work from Marie Anne O’Donnell, Swagata Basu and the editors of Nature Reviews Immunology for their work and efforts helping to run the Preprint Club. Finally, we are grateful for members of the wider scientific immunology community for their support in making the Preprint Club a resource for researchers around the world.

